# Genomic Insights into Enhanced Medium-Chain Fatty Acid Production in *Megasphaera elsdenii*

**DOI:** 10.1101/2025.08.16.670643

**Authors:** Alson Chan, Ashriel Yong, Darren Ten, Winston Koh, Yong Wei Tiong, Yee Hwee Lim, Fong Tian Wong

## Abstract

The shift toward a sustainable bioeconomy has increased interest in microbial platforms capable of producing valuable fatty acids. *Megasphaera elsdenii* is emerging as a promising candidate for fatty acid bioproduction, although its performance varies depending on the strain. This study compares two *M. elsdenii* strains, JCM 1772 (ATCC 25940) and JCM 35779, revealing that JCM 35779 produces up to 5.8 times more hexanoic acid than JCM 1772. Comparative genomic analysis identified key differences between the strains, including the loss of enzymes involved in the pyruvate-to-butyric acid pathway in JCM 35779, indicating a rerouting of carbon flux that favors chain elongation toward hexanoic acid instead of butyric acid. Overall, this work links genomic variation to metabolic phenotype, establishes JCM 35779 as a superior medium-chain fatty acid producer, and supports the development of *M. elsdenii* as an effective platform for sustainable production of medium-chain fatty acids.

**Highlights:**

- Pig-feces M. elsdenii shows >5 times higher HA production than its rumen strain.
- Genomic analysis reveals strain-specific deletions in pathways for MCFA synthesis.
- This includes etfB2, cat1 and abfD.

## 1. Introduction

In the transition toward a sustainable bioeconomy, volatile fatty acids (VFAs) are increasingly recognized as versatile platform chemicals for the production of biofuels and biochemicals. As key intermediates in biorefinery processes, VFAs derived from lignocellulosic sugars represent a promising route to reduce reliance on fossil-based resources [1]. While short-chain fatty acids (SCFAs) are already well-established industrially, medium-chain fatty acids (MCFAs), such as hexanoic acid (HA), are gaining considerable attention due to their higher energy density and value as precursors for specialty chemicals and next-generation biofuels [2,3]. Their hydrophobicity further facilitates more cost-effective downstream separation [4], increasing their appeal for large-scale biomanufacturing.

Anaerobic chain elongation via the reverse β-oxidation pathway (r-BOX) pathway has recently emerged as a central strategy for microbial MCFA production [5,6]. Within this context, *Megasphaera elsdenii* has gained interest due to its metabolic versatility and ability to utilize diverse carbon substrates, including glucose, lactate, and fermentation intermediates [6–8]. For instance, *M. elsdenii* has been shown to produce up to 4.7 g/L of HA from sucrose [7]. In comparison, *Megasphaera hexanoica*, a closely related species, has demonstrated promising but more substrate-specific MCFA production, reaching 8.9 g/L HA from lactate [9]. While *M. hexanoica* shows strong titers, *M. elsdenii* stands out with its ability to produce a broader spectrum of MCFAs, particularly BA and HA, using both glucose and lactate as feedstock [10]; albeit with variations in MCFA production profiles across strains. For example, *M. elsdenii* ATCC 25940 produced 2.6 g/L HA from 40 g/L glucose [11], whereas NCIMB 702261 and NCIMB 702410 yielded 2.2 g/L and 3.3 g/L HA, respectively, from 20 g/L sucrose [7]. In comparison, *Megasphaera* sp. MH achieved up to 4.4 g/L HA under high selective pressure (100 mM acetate) when grown on fructose [6]. Although these studies have provided valuable insights, they have primarily focused on static culture conditions and a narrow set of well-characterized strains.

Although strain-level variations in substrate utilization and metabolic efficiency have been reported among *M. elsdenii* isolates, comparative genomic analyses to elucidate these differences have not yet been conducted.Previous comparative studies suggest that significant metabolic variability exists among *M. elsdenii* strains, likely reflecting differences in genetic background and ecological origin [12–14]. This variability presents an opportunity to explore strain-specific metabolic potential in two distinct *M. elsdenii* strains exhibiting contrasting fatty acid profiles. The first species, identified as JCM 1772, originates from sheep rumen and is also annotated as ATCC 29540 [15]. The second less studied species is JCM 35779, which was recently isolated from pig feces [16]. Here, comparative genomics profiling are performed to analyse the metabolic divergence between *M. elsdenii* strains JCM 1772 and JCM 35779, with a focus on their capacity for HA production.

Overall, this study explored both production and genomic profiles, seeking to uncover the mechanisms that drive the production of MCFA, particularly on understanding the biosynthesis of BA and HA. Together, these findings open new opportunities for optimizing MCFA production, with significant potential in engineering various physiological functions and impact downstream applications in biotechnology.

## 2. Materials and methods

### 2.1 Microorganism and medium preparation

*Megasphaera elsdenii* strains, JCM 35779 and JCM 1772, were provided by the RIKEN BRC through the National BioResource Project of the MEXT, Japan. Both strains were cultivated in mYPF medium, a medium designed to support the growth of *M. elsdenii* and facilitate the production of VFAs under anaerobic conditions. The mYPF medium [6] was prepared with the following components per liter of distilled water: 10 g of yeast extract, 5 g of tryptone, 5 g of peptone, 5 g of beef extract, 10 g of fructose, 2 g of K_2_HPO_4_, 0.5 g of cysteine HCl·H_2_O, 1 mg of resazurin sodium salt, and 40 mL of a salt solution. The pH was adjusted to 6.8–7.0. The salt solution included: 0.25 g of CaCl_2_·2H_2_O, 0.5 g of MgSO_4_·7H_2_O, 1 g of K_2_HPO_4_, 1 g of KH_2_PO_4_, 10 g of NaHCO_3_, and 2 g of NaCl. Yeast extract, beef extract, and calcium chloride dihydrate were purchased from Bio Basic Asia Pacific (Singapore), Difco (USA), and Kanto Chemicals (Japan), respectively. All other chemicals were sourced from Sigma-Aldrich (USA). For routine cultivation, a Thermo Scientific™ Varioskan™ LUX plate reader (USA) was employed to monitor optical density and assess cell growth.

Genomic DNA was extracted from cell pellets using the Qiagen DNeasy Blood & Tissue Kit (Qiagen, Germany). The 16S ribosomal RNA gene was amplified by polymerase chain reaction (PCR) (see **Table S1** for primers) and the resulting amplicons were subjected to Sanger sequencing for strain identification and confirmation.

### 2.2 Fermentation

*M. elsdenii* was cultivated overnight in an anaerobic environment at 37°C with shaking at 200 rpm for 16-18 hours. The overnight culture was then used to inoculate serum bottles (50 mL, Wheaton) containing 20 mL of mYPF media. Fermentation was carried out in batch mode under anaerobic conditions, with serum bottles purged with nitrogen gas after each sample collection. Samples were routinely collected at 12-hour intervals throughout the fermentation process to measure biomass growth (via OD 600) and analyse the production of BA and HA.

### 2.3 Analytical quantification of Butyric acid (BA) and Hexanoic acid (HA)

The analytical method was adapted from Kim et al. [17] and refined through preliminary optimization. For quantification of BA and HA, 0.4 mL of culture supernatant was acidified with 15 µL of 6 M HCl, extracted with 0.1 mL of hexane, and subsequently centrifuged at 16,010 × *g* for 5 minutes. A 1 µL aliquot of the extracted hexane was then injected into a gas chromatograph (GC, Agilent 7890A, Agilent Technologies, USA) equipped with a flame ionization detector (FID) and a fused silica capillary column (HP-5, 30 m × 0.320 mm diameter × 0.5 µm film thickness). Helium was used as the carrier gas. The following GC method was modified from Volker et al. [18]. Briefly, the GC oven was initially set to 40°C with a 3-minute hold, followed by a temperature ramp to 130°C at a rate of 30°C/min, and a subsequent ramp to 300°C at 20°C/min. Concentrations of BA and HA were determined by comparing the peak areas to calibrated standard curves (**Fig. S1**).

### 2.4 Statistical Analysis

All experiments were conducted in biological triplicates. Statistical analyses were performed to evaluate differences in growth, BA and HA productions between strains and across experimental conditions. Pairwise differences among group means were assessed using Tukey test, with statistical significance defined as p < 0.05. All values are reported as mean ± standard deviation (SD).

### 2.5 Read Processing and Alignment

Raw paired-end Illumina reads were quality-filtered and adapter-trimmed using FastQC v12.1 and fastp v1.0.1 [19]. Trimmed reads were aligned to the *M. elsdenii* reference genome with BWA-MEM v0.7.19 (default parameters) [20]. Resulting SAM files were converted to BAM, sorted, and indexed using SAMtools v1.22 [21].

To evaluate gene presence and absence with nucleotide-level resolution, we leveraged annotated coding sequences (CDS) from the *M. elsdenii* reference GenBank file (Accession No: NZ_CP027570.1), extracting CDS features, including genomic coordinates, locus tags, gene names and corresponding translated protein sequences. Per-base coverage depth was calculated with SAMtools, and CDS-level proportional coverage was utilized for further gap identification and downstream gene-absence analysis.

## 3. Results and discussions

### 3.1 Comparison between JCM 35779 and JCM 1772

This study compared two *M. elsdenii* strains originating from distinct ecological niches: the well-characterized rumen-derived strain JCM 1772 (ATCC 25940), and the more recently isolated fecal strain JCM 35779, from pig feces. Notably, JCM 1772 has been extensively studied [22], and is frequently used as a benchmark strain for evaluating chain-elongating capability. In contrast, JCM 35779 remains largely unexplored, offering an opportunity to assess unexplored metabolic potential within the species.

Strain identities were confirmed through 16S rRNA phenotyping (**Table S1** and **S2**), which showed high sequence homology to reference *M. elsdenii* strains: JCM 1772 aligned with DSM 20460 (ATCC 25940), whereas JCM 35779 matched strain M215, consistent with their reported origins. Both strains were cultivated at 30°C and 37°C and to assess temperature-dependent growth behaviour. As shown in **Fig. 1A–B**, both strains exhibited markedly improved growth at 37°C, reaching higher maximal OD 600 values and achieving exponential phase more rapidly compared to cultures grown at 30°C. This temperature dependency is consistent with previous work describing *M. elsdenii* as a mesophilic organism optimized for rumen-like environments [7,12]. Across all conditions, cultures were grown in mYPF medium without pH control or in situ acid removal. As fermentation progressed, the accumulation of organic acids is known to reduce culture pH to inhibitory levels, typically pH 4.5–5.0, which adversely affects growth and metabolism in *M. elsdenii*. Thus, the growth profiles observed here reflect the uncoupled effects of acid accumulation and substrate depletion, rather than optimal fermentation performance. Notably, even under these non-optimized, acid-inhibitory conditions, the two strains displayed distinct physiological behaviours. JCM 1772 demonstrated a faster growth rate and reached a higher biomass plateau compared to JCM35779.

**Fig. 1.**
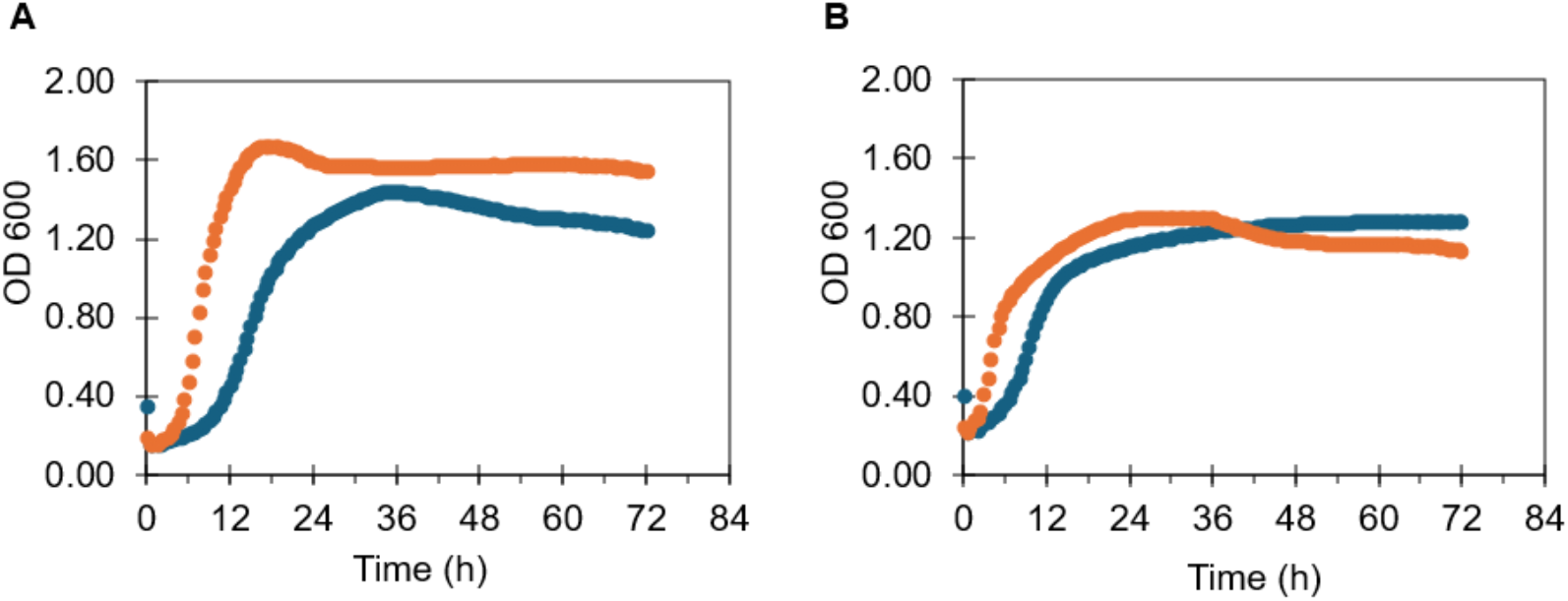
Growth profile of **(A)** JCM 1772 and **(B)** JCM 35779 at different temperatures at 30 oC (blue) and 37 oC (orange) in mYPF medium.

In addition to growth characterization, the MCFA production profiles of both strains were compared over a 48-hour batch fermentation (**Fig. 2**). Although JCM 1772 produced slightly higher levels of BA than JCM 35779, with a 1.49-fold change increase, reaching 0.18 g/L at 48 hours compared to 0.12 g/L at 48 hours for JCM 35779, the difference was not statistically significant (p>0.05), indicating that both strains maintain comparable capacities for BA production (**Fig. 2A**). In contrast, HA production showed a pronounced divergence between strains. JCM 35779 displayed a markedly superior HA-producing phenotype, achieving 1.43 g/L at 48 hours – representing a 5.78-fold increase relative to JCM 1772, which produced only 0.24 g/L (**Fig. 2B)**. This substantial difference suggests that JCM 35779 possesses a more favourable metabolic configuration for r-BOX-mediated chain elongation toward HA, corroborating the strain’s genomic signatures indicating loss of pyruvate-derived BA pathway genes and potential redistribution of carbon flux toward HA biosynthesis.

**Fig. 2.**
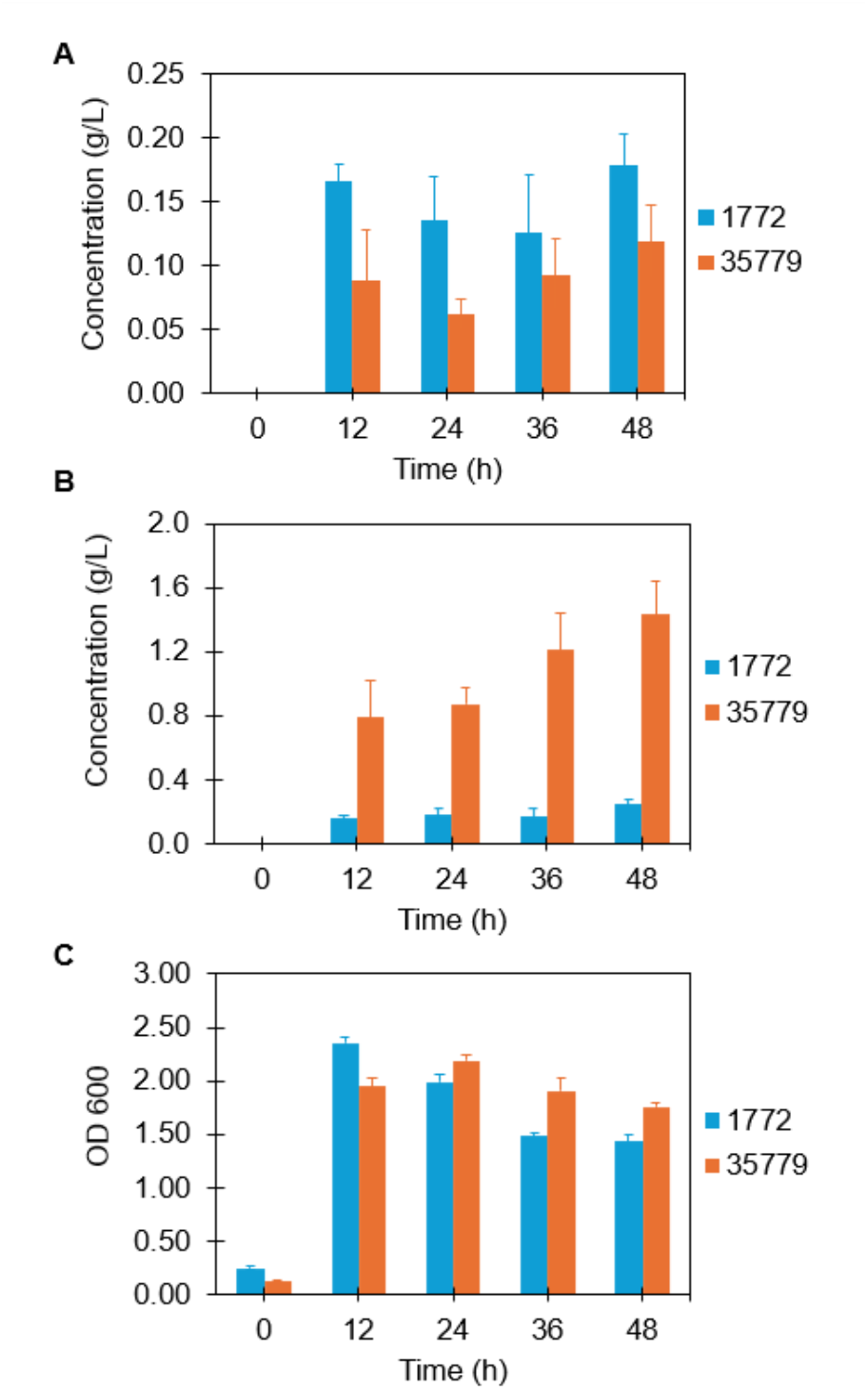
Comparison between JCM 35779 and JCM 1772 in terms of their **(A)** BA and **(B)** HA production and **(C)** growth profile (OD 600) over a period of 48 hours in mYPF medium at 37 oC. Batch fermentations were conducted in biological triplicates using 20 mL serum bottles. Statistical analysis (t-test) indicated no significant difference in BA production between the strains (p > 0.05), whereas HA production differed significantly across all time points (p<0.05).

Growth profiles revealed that JCM 1772 exhibited a slightly faster initial growth rate, reaching its maximum OD at 12 hours, whereas JCM 35779 reached its peak biomass only at 24 hours (**Fig. 2C**). However, JCM 1772 showed a noticeably sharper biomass decline following its peak, decreasing 1.64-fold by 36 h, consistent with greater sensitivity to acidification under non-pH-controlled conditions. This observation aligns with prior report that rumen-derived isolates often experience faster inhibition under acidic fermentation environments [23]. In contrast, JCM 35779 demonstrated greater physiological stability, maintaining higher OD values throughout the latter part of the fermentation. Such stability may reflect strain-specific ecological adaptation, as fecal-derived organisms are often exposed to more variable and acidic environments and therefore exhibit broader stress tolerance than rumen isolates [24].

### 3.2 Reference-Based Coverage Analysis of Genomes

Although JCM 35779 has been classified as *M. elsdenii* based on 16S rRNA analyses by Yoshikawa et al. [16] and by our analyses (**Table S2**), its genome has not been sequenced previously. In contrast, JCM 1772 is a well-established reference strain (*M. elsdenii* ATCC 25940). Despite their shared species classification, JCM 35779 exhibited markedly higher HA production (over 5-fold increase relative to JCM 1772; **Fig. 2**), suggesting the presence of substantial genetic and metabolic divergence between the strains. Such phenotypic variation among isolates of the same species is not uncommon and may arise from differences in redox metabolism, electron-donor balance, or CoA-dependent carbon flux allocation [6,25].

To systematically dissect the genetic and metabolic divergence between *M. elsdenii* strains JCM 1772 and JCM 35779, comparative genomics was employed, leveraging whole-genome resequencing and reference-based coverage analysis. Whole-genome resequencing was performed using Illumina HiSeq platform, generating paired-end reads with an average read length of 150 bp. These reads were first quality-checked and adapter-trimmed using FastQC and fastp, respectively. Subsequently, the reads were aligned to the *M. elsdenii* ATCC 25940 reference genome using the Burrows-Wheeler Aligner (BWA-MEM [20]). To quantify genomic variation, per-base coverage was computed using SAMtools. This generated a comprehensive map of nucleotide coverage across the reference genome, enabling the identification of regions with significantly reduced or absent coverage in the target strains. Discrete intervals of zero coverage were flagged as putative deletions or strain-specific absent loci, potentially representing structural variations such as insertions, deletions, or gene loss (**Fig. 3, Fig. S2**).

**Fig. 3.**
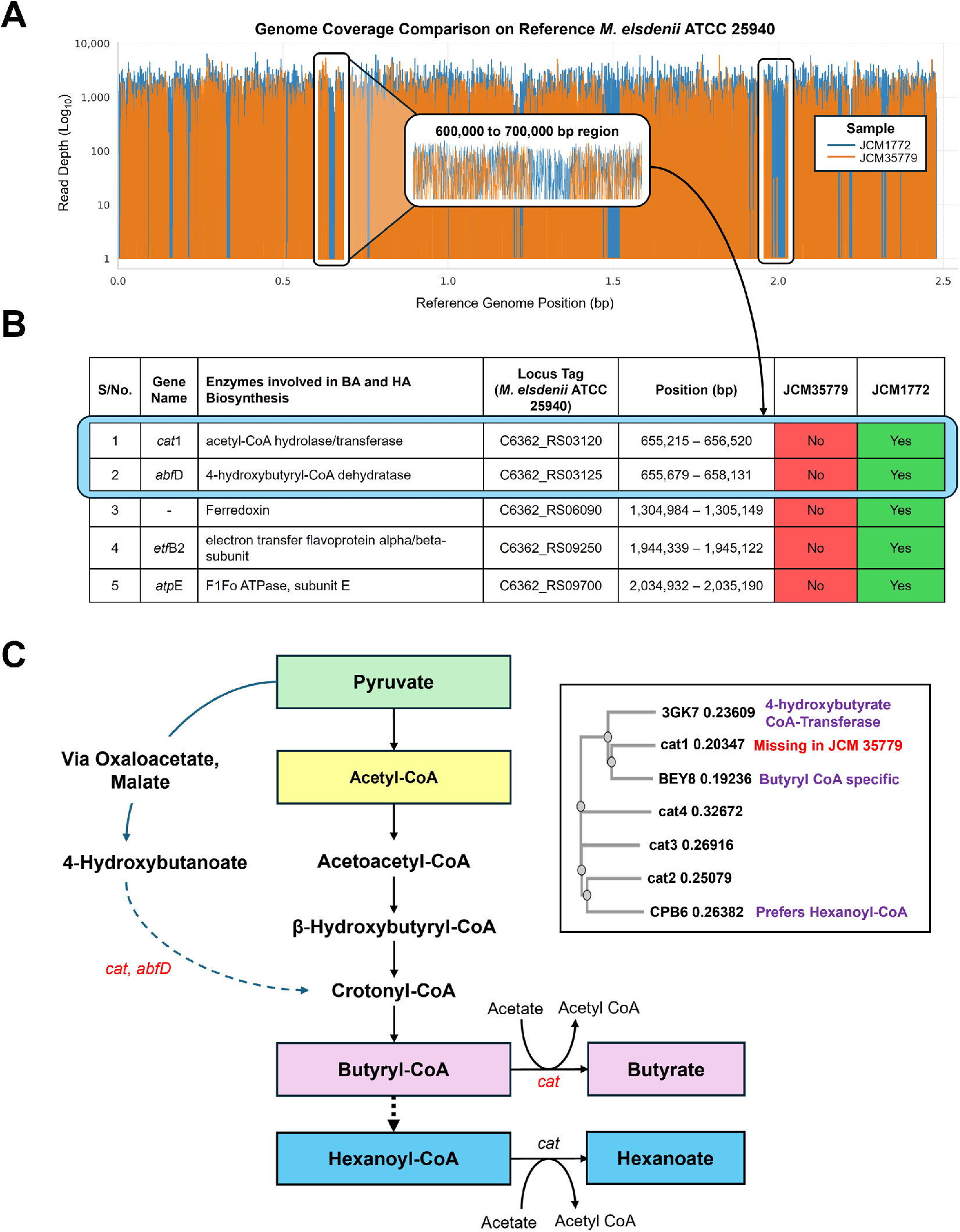
Comparative genomic analysis of *M. elsdenii* strains JCM 1772 and JCM 35779 relative to the reference genome ATCC 25940. **(A)** Read alignment coverage plots showing read alignment depth across the ATCC 25940 reference genome. JCM 1772 (blue) displays uniformly high coverage, confirming near-complete genomic similarity to the reference strain, whereas JCM 35779 (orange) exhibits multiple discrete regions of low or zero coverage, indicating putative deletions or strain-specific gene absences. **(B)** Genes associated with BA and HA biosynthesis that differ between strains. Missing genes in JCM 35779, those linked to the 4-hydroxybutyrate route (e.g., cat1, abfD), are highlighted in red. The inset phylogenetic tree compares CoA transferases from JCM 1772 with biochemically characterized enzymes (BEY8, CPB6; [26]), including the reference 4-hydroxybutyrate CoA-transferase (PDB ID: 3GK7). Additional gene annotations are provided in Tables S4–S6. **(C)** Putative biosynthesis of MCFAs, illustrating the roles of key enzymes identified in panel (B).

Per-base coverage analysis further revealed that JCM1772 and JCM35779 have varying alignment profiles with the reference genome ATCC 25940, with an alignment of 99.5 % for JCM 1772, highlighting a near identical genomic profile between JCM 1772 and the reference genome (**Table S3**). This observation confirms that JCM 1772 is correctly identified as ATCC 25940. However, JCM 35779 only has an alignment of 83.4 % with the reference genome of ATCC 25940. The coverage plot (**Fig. 3A**) revealed discrete regions of zero coverage, indicating putative deletions or strain-specific absent loci in JCM 35779.

Coordinates of no-coverage intervals were intersected with the annotated gene set of the reference genome (NCBI GenBank, *M. elsdenii* ATCC 25940, NZ_CP027570.1) to identify strain-specific gene deletions or truncations. Several missing genes were identified in JCM 35779, including those involved in BA and HA biosynthesis (**Fig. 3B, C**). Notably, this analysis revealed that JCM 35779 lacks several genes associated with short- and medium-chain fatty acid metabolism, including the β-subunit of an electron-bifurcating ETF complex (etfB2) and two enzymes involved in conversion of succinate to crotonyl-CoA, acetyl-CoA hydrolase/transferase (cat1) and 4-hydroxybutyryl-CoA dehydratase (abfD) (**Fig. 3; Table S4**). Although four CoA transferase (cat) enzymes are proposed in JCM 1772, sequence alignment with known BA/HA-specific CoA transferases [26] suggests that cat1 functions as a BA-specific CoA transferase (Figure 3B inset). cat2 is hypothesized to be similar to CPB06, which reacts with both butyryl CoA and hexanoyl CoA, with preference for hexanoyl CoA. Removing cat1 is hypothesized to not only limit crotonyl-CoA formation from 4-hydroxybutanoate but also decrease BA production, as the remaining CoA transferases show a stronger preference for hexanoyl CoA. This observation correlates with the HA/BA profiles observed in JCM 1772 and JCM 35779.

In addition, both BA and HA pathways are thermodynamically exergonic and therefore not “energy demanding,” HA synthesis requires additional chain-elongation cycles in the r-BOX pathway. Each elongation step consumes more reducing equivalents (NADH or reduced ferredoxin) than BA synthesis [27,28]. In particular, the elongation of butyryl-CoA to hexanoyl-CoA requires an extra reductive step from enoyl-CoA to acyl-CoA, increasing the overall electron demand. As such, differences in electron-bifurcating ETF complexes, ferredoxin:NAD+ oxidoreductases, and redox-linked CoA transferases between strains can alter electron-flux allocation and thereby shift the HA/BA production ratio. The absence of etfB2 and BA-associated CoA transferase genes in JCM 35779 may bias redox balance and CoA flux toward downstream elongation to HA, while the intact BA-associated gene set in JCM 1772 supports a more balanced distribution of BA and HA. These findings suggest that strain-specific redox-balancing capacity, rather than pathway energetics, underpins the distinct MCFA profiles observed between the two *M. elsdenii* strains.

## 4. Conclusion

This study analyzed both production and genomic data to understand the mechanisms behind MCFA production, specifically examining the biosynthesis pathways of BA and HA. Based on these findings, we hypothesize that genomic variations at the strain level significantly influence MCFA production in *M. elsdenii*. Comparative profiling of the rumen-derived JCM 1772 and the fecal-derived JCM 35779 revealed significant metabolic divergence despite near-identical 16S rRNA identities. Genome coverage analysis identified the loss of key r-BOX–associated genes in JCM 35779, including cat1 and abfD, which likely shifts redox and CoA flux toward chain elongation, contributing to its markedly higher HA-to-BA production ratio. These findings underscore the importance of incorporating genomic resolution into strain selection for MCFA bioprocess development.

Building on these insights, future work should integrate targeted adaptive laboratory evolution [29] or rational strain engineering with optimized fermentation strategies, such as pH control, co-substrate tuning, or in situ product removal [30], to further enhance HA productivity. Collectively, this study provides a foundation for the development of next-generation *M. elsdenii* strains and advances the broader goal of establishing robust platforms for sustainable MCFA biomanufacturing.

## Abbreviations

ALE: Adaptive Laboratory Evolution
BA: Butyric Acid
FID: Flame Ionization Detector
GC: Gas Chromatography
HA: Hexanoic Acid
HCl: Hydrochloric Acid
MCFA: Medium-Chain Fatty Acids
M. elsdenii: Megasphaera elsdenii
OD: Optical Density
OD600: Optical Density at 600 nm
SCFA: Short-Chain Fatty Acids
VFA: Volatile Fatty Acids
WT: Wild Type

## Data Availability Statement

The data supporting the findings of this study are available within the article and its supplementary materials.

## Acknowledgments

The work was financially supported by Institute of Sustainability for Chemicals, Energy and Environment (ISCE^2^), the Agency for Science, Technology and Research (A*STAR), Singapore (C233017003), and the Singapore Integrative Biosystems and Engineering Research Strategic Research & Translational Thrust (SIBER SRTT, A*STAR).

## Authors contribution

AC: Conceptualisation, methodology, validation, formal analysis, investigation, data curation, visualisation, writing (original draft). AY: Validation, investigation, writing (review & editing). YWT: Validation, investigation, formal analysis, visualisation, writing (review & editing). YHL: Conceptualisation, writing (original draft), resources, supervision, project administration, funding acquisition, writing (review & editing). FTW: Conceptualisation, writing (original draft), investigation, resources, formal analysis, supervision, project administration, funding acquisition, writing (review & editing). D.T.: Validation, investigation, writing (review & editing). W.K.: Validation, investigation, supervision, writing (review & editing).

## Data availability

The authors declare that the data supporting the findings of this study are available within the paper and its Supplementary Information files. Should any raw data files be needed they are available from the corresponding author upon reasonable request.

## Declarations

### Ethics approval and consent to participate

Not applicable.

### Consent for publication

Not applicable.

## References

1. Kobayashi H, Fukuoka A. Synthesis and utilisation of sugar compounds derived from lignocellulosic biomass. Green Chemistry [Internet]. 2013;15:1740–63. Available from: 10.1039/C3GC00060E

2. Stamatopoulou P, Malkowski J, Conrado L, Brown K, Scarborough M. Fermentation of organic residues to beneficial chemicals: A review of medium-chain fatty acid production. Processes [Internet]. 2020;8:1–25. Available from: 10.3390/pr8121571

3. Wu Q, Jiang Y, Chen Y, Liu M, Bao X, Guo W. Opportunities and challenges in microbial medium chain fatty acids production from waste biomass. Bioresource Technology [Internet]. 2021;340:125633. Available from: 10.1016/j.biortech.2021.125633

4. Ho Ahn J, Hwan Jung K, Seok Lim E, Min Kim S, Ok Han S, Um Y. Recent advances in microbial production of medium chain fatty acid from renewable carbon resources: A comprehensive review. Bioresource Technology [Internet]. 2023;381:129147. Available from: 10.1016/j.biortech.2023.129147

5. Liu Y, Chen L, Duan Y, Li R, Yang Z, Liu S, et al. Recent progress and prospects for chain elongation of transforming biomass waste into medium-chain fatty acids. Chemosphere [Internet]. 2024;355:141823. Available from: 10.1016/j.chemosphere.2024.141823

6. Jeon BS, Choi O, Um Y, Sang BI. Production of medium-chain carboxylic acids by Megasphaera sp. MH with supplemental electron acceptors. Biotechnology for Biofuels [Internet]. 2016;9:1–9. Available from: 10.1186/s13068-016-0549-3

7. Choi K, Jeon BS, Kim BC, Oh MK, Um Y, Sang BI. In situ biphasic extractive fermentation for hexanoic acid production from sucrose by megasphaera elsdenii NCIMB 702410. Applied Biochemistry and Biotechnology. 2013;171:1094–107.

8. Sträuber H, Bühligen F, Kleinsteuber S, Dittrich-Zechendorf MD-Z. Carboxylic acid production from ensiled crops in anaerobic solid-state fermentation - trace elements as pH controlling agents support microbial chain elongation with lactic acid. Engineering in Life Sciences. 2018;18:447–58.

9. Kang S, Kim H, Jeon BS, Choi O, Sang BI. Chain elongation process for caproate production using lactate as electron donor in Megasphaera hexanoica. Bioresource Technology [Internet]. 2022;346:126660. Available from: 10.1016/j.biortech.2021.126660

10. Gildemyn S, Molitor B, Usack JG, Nguyen M, Rabaey K, Angenent LT. Upgrading syngas fermentation effluent using Clostridium kluyveri in a continuous fermentation. Biotechnology for Biofuels. 2017;10:1–15.

11. Roddick FA, Britz ML. Production of hexanoic acid by free and immobilised cells of Megasphaera elsdenii: Influence of in-situ product removal using ion exchange resin. Journal of Chemical Technology and Biotechnology. 1997;69:383–91.

12. Nelson RS, Peterson DJ, Karp EM, Beckham GT, Salvachúa D. Mixed carboxylic acid production by megasphaera elsdenii from glucose and lignocellulosic hydrolysate. Fermentation. 2017;3.

13. Cabral L da S, Weimer PJ. Megasphaera elsdenii: Its Role in Ruminant Nutrition and Its Potential Industrial Application for Organic Acid Biosynthesis. Microorganisms. 2024;12:1–27.

14. Lee NR, Lee CH, Lee DY, Park JB. Genome-scale metabolic network reconstruction and in silico analysis of hexanoic acid producing megasphaera elsdenii. Microorganisms. 2020;8.

15. Elsden SR, Volcani BE, Gilchrist MC, Lewis D. Properties of a fatty acid forming organism isolated from the rumen of sheep. Journal of Bacteriology. 1956;72:681–9.

16. Yoshikawa S, Itaya K, Hoshina R, Tashiro Y, Suda W, Cho Y, et al. Thermophile-fermented feed modulates the gut microbiota related to lactate metabolism in pigs. Journal of applied microbiology. 2024;135.

17. Kim H, Jeon BS, Sang BI. An Efficient New Process for the Selective Production of Odd-Chain Carboxylic Acids by Simple Carbon Elongation Using Megasphaera hexanoica. Scientific Reports [Internet]. 2019;9:1–10. Available from: http://dx.doi.org/10.1038/s41598-019-48591-6

18. Volker AR, Gogerty DS, Bartholomay C, Hennen-bierwagen T, Zhu H, Bobik TA. Fermentative production of short-chain fatty acids in Escherichia coli. 2014;1513–22.

19. Chen S, Zhou Y, Chen Y, Gu J. fastp: an ultra-fast all-in-one FASTQ preprocessor. Bioinformatics. 2018;34:884–90.

20. Li H, Durbin R. Fast and accurate short read alignment with Burrows – Wheeler transform. 2009;25:1754–60.

21. Li H, Handsaker B, Wysoker A, Fennell T, Ruan J, Homer N, et al. The Sequence Alignment/Map format and SAMtools. Bioinformatics. 2009;25:2078–9.

22. Hatmaker EA, Klingeman DM, Dell KBO, Riley LA, Papanek B. Complete Genome Sequences of Two Megasphaera elsdenii, NCIMB 702410 and ATCC 25940. Microbiology Resource Announcements. 2019;8:1–2.

23. Candry P, Huang S, Carvajal-arroyo JM, Rabaey K, Ganigue R. Enrichment and characterisation of ethanol chain elongating communities from natural and engineered environments. 2020;1–10.

24. Donaldson GP, Lee SM, Mazmanian SK. Gut biogeography of the bacterial microbiota. Nature Reviews Microbiology. 2016;14:20–32.

25. Jarboe LR, Royce LA, Liu P. Understanding biocatalyst inhibition by carboxylic acids. Frontiers in Microbiology. 2013;4:1–8.

26. Yang Q, Guo S, Lu Q, Tao Y, Zheng D, Zhou Q, et al. Butyryl/Caproyl-CoA:Acetate CoA-transferase: cloning, expression and characterization of the key enzyme involved in medium-chain fatty acid biosynthesis. Bioscience Reports. 2021;41:1–14.

27. Angenent LT, Richter H, Buckel W, Spirito CM, Steinbusch KJJ, Plugge CM, et al. Chain Elongation with Reactor Microbiomes: Open-Culture Biotechnology To Produce Biochemicals. Environmental Science & Technology. 2016;50:2796–810.

28. Zhao J, Ma H, Gao M, Qian D, Wang Q, Shiung S. Advancements in medium chain fatty acids production through chain elongation : Key mechanisms and innovative solutions for overcoming rate-limiting steps. Bioresource Technology. 2024;408:131133.

29. Lakhssassi N, Baharlouei A, Meksem J, Hamilton-Brehm SD, Lightfoot DA, Meksem K, et al. Ems-induced mutagenesis of clostridium carboxidivorans for increased atmospheric CO2 reduction efficiency and solvent production. Microorganisms. 2020;8:1–13.

30. Mcdowall SC, Braune M, Nitzsche R. Recovery of bio-based medium-chain fatty acids with membrane filtration. Separation and Purification Technology. 2022;286.

