## Supplemental figures and tables for "Genomic Insights into Enhanced Medium-Chain Fatty Acid Production in *Megasphaera elsdenii*"

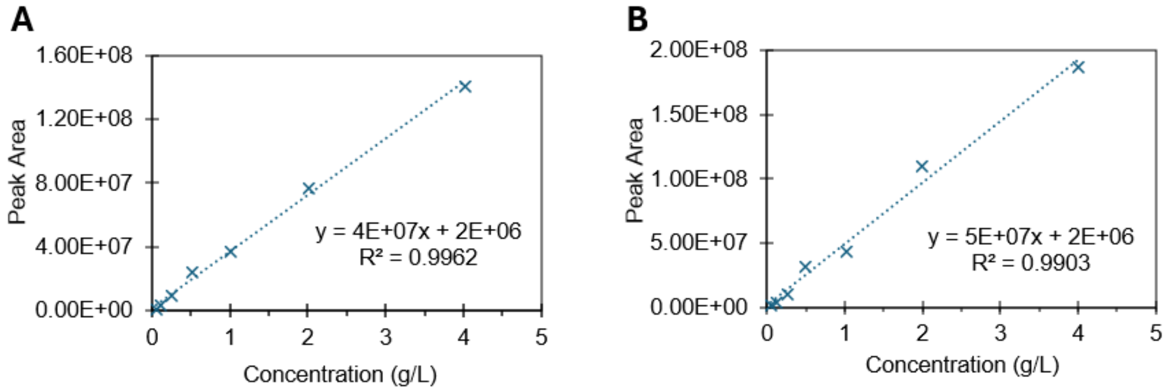


**Fig. S1.** Calibration curves for the quantification of **(A)** BA, and **(B)** HA production.

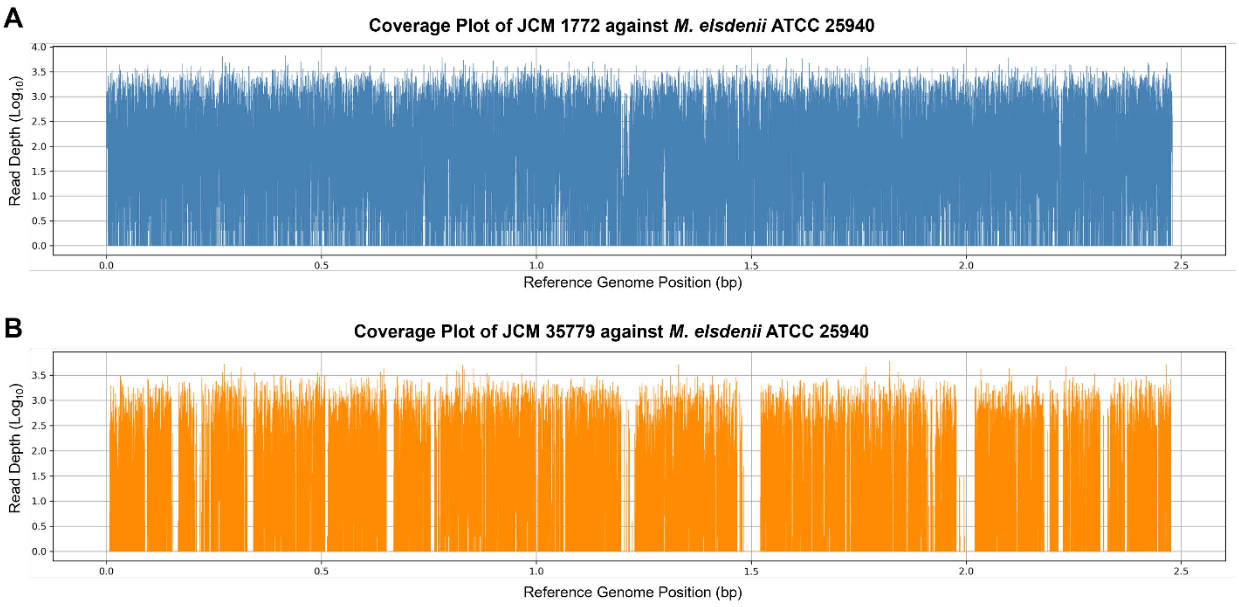


**Fig. S2.** Alignment coverage plots of **(A)** JCM 1772 and **(B)** JCM 35779 against *M. elsdenii* ATCC 25940.

### Tables

**Table S1.** Primers used for 16S rRNA sequencing.

| Primer | Sequence (5' → 3') | References |
|---|---|---|
| Mega-142F | GATGGGGACAACAGCTGGA | Development of a 16S rRNA Gene Primer and PCR-Restriction Fragment Length Polymorphism Method for Rapid Detection of Members of the Genus Megasphaera and Species-Level Identification - PMC |
| Mega-X | GACTCTGTTTTTGGGGTTT | |

**Table S2.** 16S rRNA homology results for *JCM 1772* and *JCM 35779* from BLAST analysis.

| Strain ID | Description (database) | Description | Query Cover | Per. Ident |
|---|---|---|---|---|
| JCM 1772 Megasphaera elsdenii | Megasphaera elsdenii gene for 16S ribosomal RNA, partial sequence, strain: JCM 1772 | Megasphaera elsdenii DSM 20460 strain ATCC 25940 chromosome, complete genome | 99% | 97.49% |
| JCM 35779 Megasphaera elsdenii | | Megasphaera elsdenii M215 gene for 16S ribosomal RNA, partial sequence | 99% | 99.56% |

**Table. S3.** Sequencing reads and alignment coverage of JCM 1772 and JCM 35779 against *M. elsdenii* ATCC 25940.

| | **JCM 1772** | **JCM 35779** |
|---|---|---|
| **Total reads (150-bp)** | 8,667,618 | 4,805,784 |
| **Primary reads** | 8,667,502 | 4,792,820 |
| **Secondary alignments** | 116 | 12,964 |
| **Total reads mapped to the reference genome** | 8,654,096 (99.84%) | 4,026,861 (83.79%) |
| **Total reads mapped to the reference genome (correct orientation)** | 8,625,292 (99.51%) | 3,998,198 (83.42%) |
| **Singletons** | 4514 | 3037 |

**Table. S4.** MCFA proteins exclusively to JCM 1772 but absent in JCM 35779.

| S/No. | Annotated CDS from Reference Genome | Locus Tag | Start | End | CDS Length | Amino Acid Length |
|---|---|---|---|---|---|---|
| 1 | Acetyl-CoA hydrolase/transferase family protein | C6362_RS03120 | 655,215 | 656,520 | 1305 | 434 |
| 2 | *MaoC* family dehydratase | C6362_RS03135 | 659,016 | 659,748 | 732 | 243 |
| 3 | Radical SAM protein | C6362_RS09115 | 1,913,528 | 1,914,407 | 879 | 292 |
| 4 | 4Fe-4S cluster-binding domain-containing protein | C6362_RS09500 | 1,990,792 | 1,991,332 | 540 | 179 |
| 5 | NAD(P)H-dependent oxidoreductase | C6362_RS10975 | 2,321,974 | 2,322,685 | 711 | 236 |

**Table S5.** Distant Homologs of absent MCFA-associated Proteins in JCM35779.

| S/No. | Query MCFA Proteins | Query Locus ID | BLASTp Hit Proteins | Hit Locus ID | E-value | Identity (%) | CDS-level Coverage in JCM 35779 (%) |
|---|---|---|---|---|---|---|---|
| 1 | Acetyl-CoA hydrolase/transferase family protein | C6362_RS03120 | Acetyl-CoA hydrolase/transferase family protein | C6362_RS03615 | $1.22\times10^{-92}$ | 40 | 98.52 |
| 2 | Acetyl-CoA hydrolase/transferase family protein | C6362_RS03120 | Acetyl-CoA hydrolase/transferase family protein | C6362_RS03540 | $2.8\times10^{-89}$ | 36.78 | 77.46 |
| 3 | Acetyl-CoA hydrolase/transferase family protein | C6362_RS03120 | Acetyl-CoA hydrolase/transferase C-terminal domain-containing protein | C6362_RS03660 | $4.76\times10^{-67}$ | 37.87 | 29.79 |

| | | | | | | | |
|---|---|---|---|---|---|---|---|
| 4 | Radical SAM protein | C6362_RS 09115 | Radical SAM protein | C6362_RS 02945 | $9.52\times 10^{-53}$ | 37.31 | 70.16 |
| 5 | Radical SAM protein | C6362_RS 09115 | elongator complex protein 3 | C6362_RS 01675 | $7.41\times 10^{-6}$ | 24.06 | 96.88 |
| 6 | Radical SAM protein | C6362_RS 09115 | radical SAM family heme chaperone HemW | C6362_RS 09630 | $4.73\times 10^{-5}$ | 22.17 | 95.38 |
| 7 | Radical SAM protein | C6362_RS 09115 | radical SAM protein | C6362_RS 02340 | $2.6\times 10^{-4}$ | 24.47 | 91.44 |
| 8 | Radical SAM protein | C6362_RS 09115 | TIGR03960 family B12-binding radical SAM protein | C6362_RS 11555 | $9.1\times 10^{-4}$ | 29.63 | 90.42 |

To determine putative homologs of selected MCFA-associated proteins, intra-genomic BLASTp searches were performed against the reference genome of *M. esldenii* ATCC 25940. Protein sequences of interest, along with their corresponding locus tags, were extracted from the annotated GenBank file and used as queries against a custom BLASTP database constructed from the reference proteome. BLASTp was executed with an E-value threshold of $1\times 10^{-5}$ and a minimum amino acid identity of 20%, to accommodate distant homologs. Previously computed per-base CDS coverage was cross-referenced to evaluate the presence of homologous in JCM 35779. No homologous hits were found for *MaoC* family dehydratase, 4Fe-4S cluster-binding domain-containing protein and NAD(P)H-dependent oxidoreductase.
